# IPCO: Inference of Pathways from Co-variance analysis

**DOI:** 10.1101/686923

**Authors:** Mrinmoy Das, Tarini Shankar Ghosh, Ian B. Jeffery

## Abstract

Key aspects of microbiome research are accurate identification of taxa followed by the profiling of their functionality. Amplicon profiling based on the 16S ribosomal DNA sequence is a ubiquitous technique to identify and profile the abundances of the various taxa. However, it does not provide information on their encoded functionality. Predictive tools which can accurately extrapolate the functional information of a microbiome based on taxonomic profile composition is essential. At present the applicability of these tools is however limited due to requirement of reference genomes from known species. We present IPCO (Inference of Pathways from Co-variance analysis), a new method of inferring functionality for 16S-based microbiome profiles independent of reference genomes. IPCO utilises the biological co-variance observed between paired taxonomic and functional profiles and co-varies it with the queried dataset. It outperforms other established methods both in terms of sample and feature profile prediction. Validation results confirmed that IPCO can replicate observed biological signals seen within shotgun and metabolite profiles. Comparative analysis of predicted functionality profiles with other popular 16S-based functional prediction tools indicates significantly lower performance with predicted functionality showing little to no correlation with paired shotgun features across samples. IPCO is implemented in R and available from https://github.com/IPCO-Rlibrary/IPCO.

## Introduction

Microbiome research has expanded exponentially over the years. Microbial communities have significant roles in maintaining health and the environment as well as their roles food and industrial processes is being recognised to be very important [1, 2]. Study of microbiome communities fundamentally falls under two categories: the taxonomic compositions determined by amplicon sequencing (16S or marker gene) or metagenomic whole genome shotgun sequencing (mWGS) and their functional capabilities which requires mWGS to identify genes and pathways. Despite the reducing cost of mWGS, amplicon sequencing still remains popular due to its relatively low cost, quicker computation time, less data size, and ability to detect rare organisms due to its greater sampling depth compared to mWGS. Comparison of these two approaches have discussed both the advantages and disadvantages of these methods [3, 4]. However, amplicon sequencing is limited to providing only taxonomic information of the microbial communities.

A number of tools can predict the functional potential of the microbial communities obtained from 16S sequencing, the most cited being PICRUSt which was published as early as 2013 [5]. Other widely used tools that were developed later are Tax4Fun (2015) [6], and Piphillin (2016) [7]. All of these tools rely on functionally annotated reference genomes. The difference between them is the approach used to assign functional annotation to those references whose functionality is unknown and that used to map the amplicon data to these references. PICRUSt works by considering the phylogenetic tree and the distance to the closest functionally annotated reference microbe [5]. It relies on GreenGenes database [8] for matching the references to the queried amplicon data. However, the major limitation of PICRUSt comes forth in the case of microbes which do not have sequenced/annotated genes of phylogenetically close relatives in the reference database. Tax4Fun and Piphillin implements BLAST and global alignment respectively between the amplicon data and the reference genomes obtained from different databases [6, 7]. All of these tools uses KEGG orthologs (KOs) [9] annotation from reference genomes to predict the functionality by computing against the amplicon abundance. However, the limitation of all these methods is the requirement of sequenced/annotated referenced genomes.

To overcome constrains of limited reference taxa and lack of feature correlation across samples, an alternative approach would be to co-vary the taxonomic abundance and functionality of reference dataset and associate the covariance trends with the taxonomic abundance of the queried dataset. In this paper, we present IPCO, a tool based on a novel approach for inferring functionality to a 16S dataset. The primary feature of our method is that it does not depend on the presence of sequenced and annotated genomes directly. IPCO is an application of double co-inertia analysis involving the RLQ method (R-mode; Q-mode; and L-link between R and Q) [10, 11] between a paired taxonomic and its functional dataset and the 16S dataset for which functionality to be inferred, to assign a value to the functional profiles for the samples of a 16S dataset. Co-inertia analysis measures the concordance between two datasets, and maximises the squared covariance projected by two datasets [12, 13]. In a paired taxa and functional profile datasets one would expect that alterations in the taxonomic profiles naturally should also reflect changes in its functional potential. Co-inertia is further extended by application of RLQ method, which integrates a third dataset (amplicon dataset in this case) which analyses co-inertia of the three datasets simultaneously and obtains a set of scores for the functional dataset and the amplicon dataset weighted by the paired taxonomic dataset.

IPCO’s performance was compared with PICRUSt, Tax4Fun and Piphillin in terms of both samples and features correlation with KEGG pathways. IPCO can associate MetaCyc pathway profiles also which is compared against paired mWGS data only as published tools currently work only on KEGG. Correlation of mWGS functional profile against its paired bile acids and short chain fatty acids (SCFAs) metabolite profile confirmed the biological signals associated with the different pathways. Using functionality inferred from IPCO, these biological signals were reproduced and showed consistency as observed with mWGS dataset.

## Methodology

### IPCO algorithm

IPCO is an implementation of the RLQ analysis which is also known as fourth corner analysis. It requires a reference taxonomic and functional paired dataset along with a 3^rd^ dataset which is the 16S dataset for which the functional potential need to be inferred. RLQ analysis a double co-inertia method which explores two datasets (R and Q) through a mediator dataset (L). IPCO implements RLQ to associate the functional profiles (R) with a 16S profile dataset (Q) which is the 16S dataset for which functions need to be inferred through a mediator taxonomic profile dataset (L). R and L datasets are related as they have the same samples (paired) and are the reference datasets. Q and L datasets are related as they have the same taxa. Functional profiles of R and taxonomic profiles of Q are standardised and scaled through the weighted average where the weights of the samples and taxa are obtained from L dataset. Through RLQ methodology, we obtain a R’LQ product table which an association matrix of between R and Q medicated through L abundance. In IPCO, we re-standardise the R’LQ products by adding the weighted average of the functional potential back to the association matrix to obtain inferred functional profiles for the samples of Q dataset.

### Data collection

In the current study, human microbiome taxonomic and functional profile dataset were obtained from the HMP project [14, 15] and curatedMetagenomicData R library [16]. mWGS functional (UniRef genefamilies) and taxonomic profiles datasets were obtained using the R library curatedMetagenomicData and paired V3-V5 16S rRNA OTU table was obtained from 16SHMPData R library [17]. Paired datasets were obtained for nasal, oral(buccal cavity), skin and stool samples. Representative OTUs of V3-V5 regions were downloaded from the HMP website [14, 15].

A larger reference dataset consisting of functional and taxonomic profiles generated from only mWGS data were also obtained from the curatadMetagenomicData. This set is comprised of 1180 healthy samples from various cohort as describes in Ghosh et al (manuscript in preparation).

Paired 16S and mWGS of the environmental dataset are described in Tessler *et al.* [18] and downloaded from the NCBI SRA (PRJNA389803, PRJNA310230). The environmental 16S rRNA and mWGS data was quality filtered using trimmometric (v.0.38) [19]. The quality filtered 16S rRNA sequences were processed as described in Tran *et al.* [20] to obtain the OTU table. The quality filtered mWGS data was processed using HUMAnN2 [21] to obtain mWGS derived taxonomic and functional profiles.

MetaCyc and KEGG pathway mapping files provided with HUMAnN2 and HUMAnN1 were filtered to remove all known eukaryotic pathways. All samples from all datasets were processed against the filtered MetaCyc mapping file to obtain the MetaCyc pathway abundance and coverage datasets.

The UniRef genefamilies dataset for all the samples were regrouped to KEGG Orthologs IDs using humann2_regroup_table.py script and the KEGG to UniRef mapping provided in HUMAnN2 utilities. The regrouped KOs were processed using HUMAnN2 and the filtered HUMAnN1 KEGG pathways legacy database to obtained KEGG pathway profiles for the mWGS dataset.

All OTU datasets were filtered to remove samples with a sequencing depth of less than 1000 reads. Samples removed from OTU datasets were omitted from their paired mWGS datasets also. Functional information stratified with taxonomic labels were also removed from the functional datasets. UNMAPPED, UNGROUPED and UNINTEGRATED were removed after normalising the datasets.

### Data normalisation and transformations

Taxonomic and functional abundance datasets were transformed using the following transformations: Z-scaling, proportion normalisation, log10 on rarefied and log10 on proportional data with 1e-05 added as minimum count value, Hellinger transformation [22] and centred log ratio transformation [23]. These transformations were investigated to identify the transformation best suited for IPCO.

### Validation of IPCO predictions

As IPCO is independent of reference genomes and dependent on reference datasets, we implemented a bootstrap strategy to evaluate its predictions (**Figure 1**). A subset of the samples from the 16S table were randomly selected and considered as table Q. The samples omitted in Q formed the taxonomic table L and were matched with its pathway abundance dataset to obtain a paired functional (R) and taxonomy (L) datasets thus removing pathways information for the samples present in table Q. Using IPCO on R, L and Q table, pathway profiles were obtained for the samples from Q. Both inferred sample and features (pathways) values were correlated using Spearman correlation with the mWGS pathway dataset for those samples present in Q. The correlations values asserted how close the sample and feature values were to the actual profiles. It was repeated with 100 iterations to randomly subsample different samples each time. An average was taken for both samples to sample and pathways to pathways correlation values from the 100 iterations.

**Figure 1.**
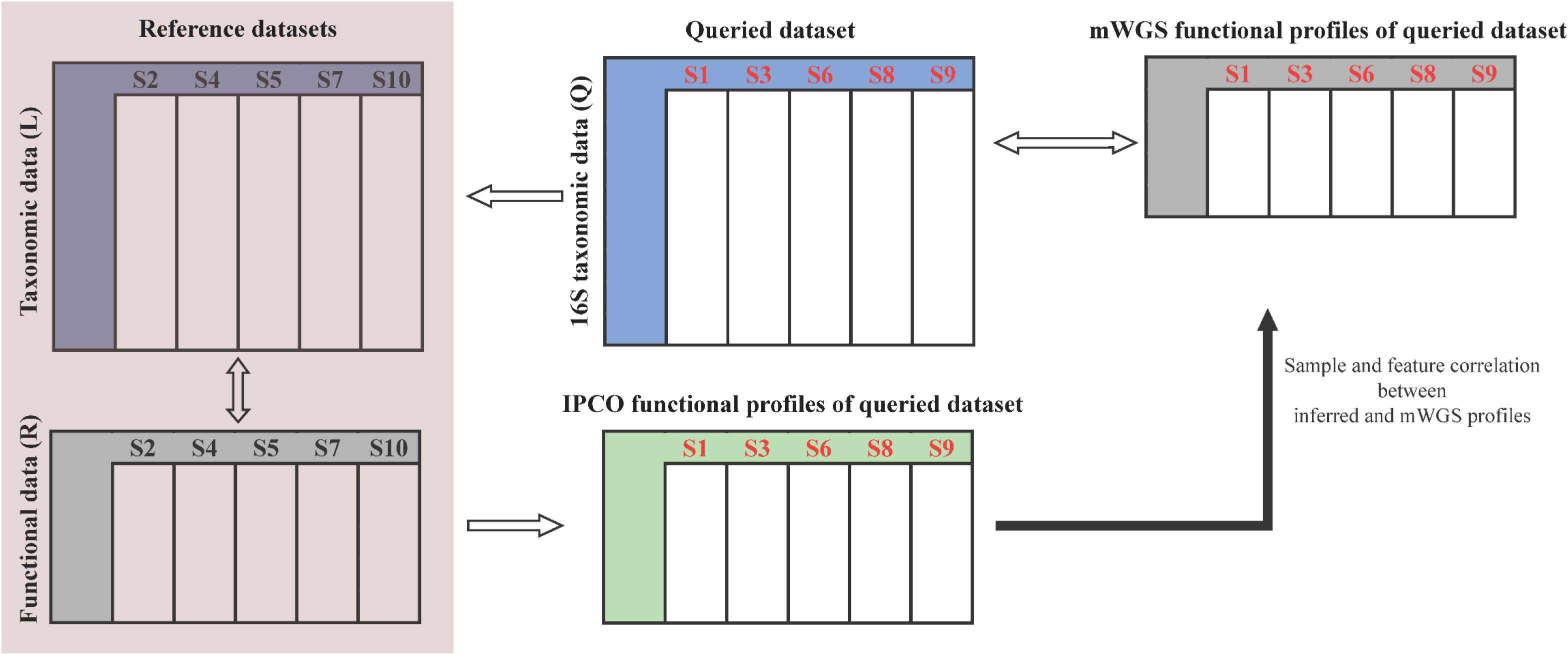
Implementation of IPCO and bootstrap iterations The datasets were randomly subsampled into query and reference dataset. Reference dataset consist of taxonomic and functional profiles for the same samples. Reference taxonomic and queried dataset consist of different samples but are mapped to same taxa. The inferred profiles is correlated against the mWGS functional profiles for the query dataset to obtain the degree of associations. 100 bootstrap iteration were carried out to randomly generate different subsamples and measure the degree of association for each iteration

### Thresholds and filtering

As IPCO is based on covariance between the datasets, it was implemented at various taxonomic levels and various subsampling threshold levels to retain a defined proportions of samples in the reference (10%, 30%, 50%, 70% and 80%) and investigate the best suited sample size and taxonomic level.

Taxonomy was assigned to the representative sequences using the RDP database (v.11.4) [24] implemented in mothur [25]. In addition, SPINGO [26] was used for species assignment using the RDP database (v.11.2). All levels of taxonomic classification were classified to a threshold of ≥ 80% confidence. At any level, if the classified threshold was below 80%, it was set as unclassified.

### Functional predictions using other methods

For PICRUSt, the representative 16S rRNA sequences were mapped to green-genes v13.5 to obtained closed representative OTU table and processed through the PICRUSt pipeline to obtain KO profiles. The KEGG pathways abundance were obtained using HUMAnN2 by processing the KO profiles with filtered HUMAnN1 KEGG legacy database.

For Tax4Fun, the taxonomic information were assigned to the representative sequences using SILVA123 downloaded from Tax4Fun website. The OTU table with taxonomic profiles were processed through Tax4Fun R library to obtain KO profiles and KEGG pathway profiles. All processing was carried out with default settings.

For Piphillin, OTU dataset and the representative OTU sequences were formatted as per requirement for Piphillin and all files were uploaded to the Piphillin website. An identity threshold of 90% was used and the KEGG database October 2018 version was used. Using 97% returned very low number of hits to reference genomes.

Sample and KEGG pathways abundance correlations were calculated by comparing the predicted samples and KEGG pathways values obtained by the three tools against the mWGS generated profiles using Spearman correlation. KEGG pathways present in both predicted and mWGS dataset were considered.

### Validation on independent dataset

To validate the results on an external independent dataset, two steps of analysis were carried out. All tests were implemented on an in-house elderly community dataset [27] which contains the mWGS, 16S and metabolome datasets for the same samples.

Firstly, it was investigated whether using mWGS species level dataset is sufficient in reference set (table L) to generate inferred functionality for the independent dataset. If this generates comparable results as 16S taxonomic reference, then it would allow generation of a larger reference R and L table by adding more samples from curatedMetagenomicData hub which would incorporate more functionality else would have to use HMP or similar 16S dataset as table L. Species level dataset were obtained for the 16S elderly and HMP 16S as described earlier. IPCO was implemented with the R table as HMP pathways dataset, L table as HMP closed OTU or 16S species or HMP mWGS species dataset and Q table as elderly 16S closed OTU or species dataset. Samples and common features from inferred table were correlated with the paired elder mWGS functional dataset.

The species and functional dataset for the 1180 healthy samples were obtained from curatedMetagenomicData. IPCO was implemented with R table as reference pathways, L table as species mWGS dataset and Q table as elderly species 16S dataset. The inferred functionality dataset for the elderly was correlated with the elderly mWGS functionality dataset to obtain samples and features correlation. Correlations obtained from using this healthy reference were compared with the correlation values from using HMP as reference.

After this, to investigate the biological signal in the predicted functionality datasets, functional profiles were obtained using PICRUSt, Tax4Fun and Piphillin from the elderly community 16S dataset. Two types of metabolites bile acids and short chain fatty acids (SCFAs) which are widely studied in microbiome research were considered for investigation. The metabolite dataset was log10 transformed on the measured metabolite level after adding 1e-05 as minimum count value. The mWGS functional profiles were correlated with the metabolite profiles and the directionality, degree of association and significance was noted. The inferred functions obtained from all the tools including IPCO were then correlated with the same metabolites and the results were compared with mWGS results to investigate the direction, correlation and significance (p-value ≤ 0.05). Only key pathways responsible for these metabolites were considered and agreement with mWGS results in terms of directionality and significance was considered to be correct results and change in directionality or significance was considered as false results.

### Statistical analysis

All analysis was carried out in R (v.3.5.1) [28]. All correlations measured were carried out using Spearman correlation. Kruskal-Wallis test was used as applicable. Dunn’s test using dunn.test library (v.1.3.5) [29] was used for pairwise comparison at different taxa and sample threshold levels. P-value adjustment was carried out using Benjamini-Hochberg procedure. Covariance between paired taxonomic and functional dataset was investigated with co-inertia analysis using ade4 (v.1.7.13) library [30]. Significance of co-inertia was determined by with randtest. Plots were created using ggplot2 (v.3.1.0) [31], RColorBrewer (v.1.1.2) [32] and gridExtra (v.2.2.1) [33] libraries.

## Results

### Description of the study cohort

In the current study, **table 1** describes the cohort retained after removing samples with low sequencing depths and stratified functional features for the initial analysis. The samples retained were investigated using the IPCO, PICRUSt and Tax4Fun to evaluate the performance of the tools.

**Table 1.** Number of mWGS features and samples retained in the initial analysis

|  | Samples | KEGG pathways | MetaCyc | OTUs |
| --- | --- | --- | --- | --- |
| HMP nasal | 61 | 129 | 659 | 7464 |
| HMP oral | 71 | 133 | 625 | 13696 |
| HMP skin | 8 | 132 | 593 | 3492 |
| HMP stool | 87 | 128 | 712 | 9966 |
| Environmental | 37 | 114 | 497 | 1185 |

Total number of samples and features used for the investigating the predictability of IPCO from different sites and datasets.

IPCO was initially implemented in the HMP stool cohort to investigate the effects of different data transformation and the taxonomic level and sample size threshold.

### Data transformation of the reference datasets affects IPCO results

In IPCO, the covariance between the three dataset (R, L and Q) was observed to vary depending on the transformation and normalisation method used. Investigation with the transformation/normalisation mentioned in the methods is shown in **supplementary figure 1** and **Supplementary table 1**. The Hellinger transformation was observed to be best suited as it had a higher RV coefficient for both samples and features and similar correlation values compared to other methods. Observed RV coefficients were significant (p-value ≤ 0.05) for all cases. Hence this transformation is implemented as default for all further analysis.

### Lowest taxonomic levels and high samples size provides best results with IPCO

Implementation of IPCO with 100 bootstraps for subsampling at different reference sample sizes on both KEGG and MetaCyc pathway abundance datasets showed that the best sample and feature correlation were observed with the lowest taxonomic levels and highest sample size (**Figure 2A-B**, **supplementary figure 2A-B**). No significant difference was observed in the sample correlation for any reference dataset size expect for between 10% and other reference sizes at family level only in the MetaCyc dataset **(Supplementary table 2)**. However, the feature correlation increased with increase in sample size and at lowest taxonomy levels. No significant difference was observed for feature correlation using a reference size of at least 30% or larger in KEGG pathways and 50% or more for MetaCyc at different taxonomic levels **(Supplementary table 2)**.

**Figure 2.**
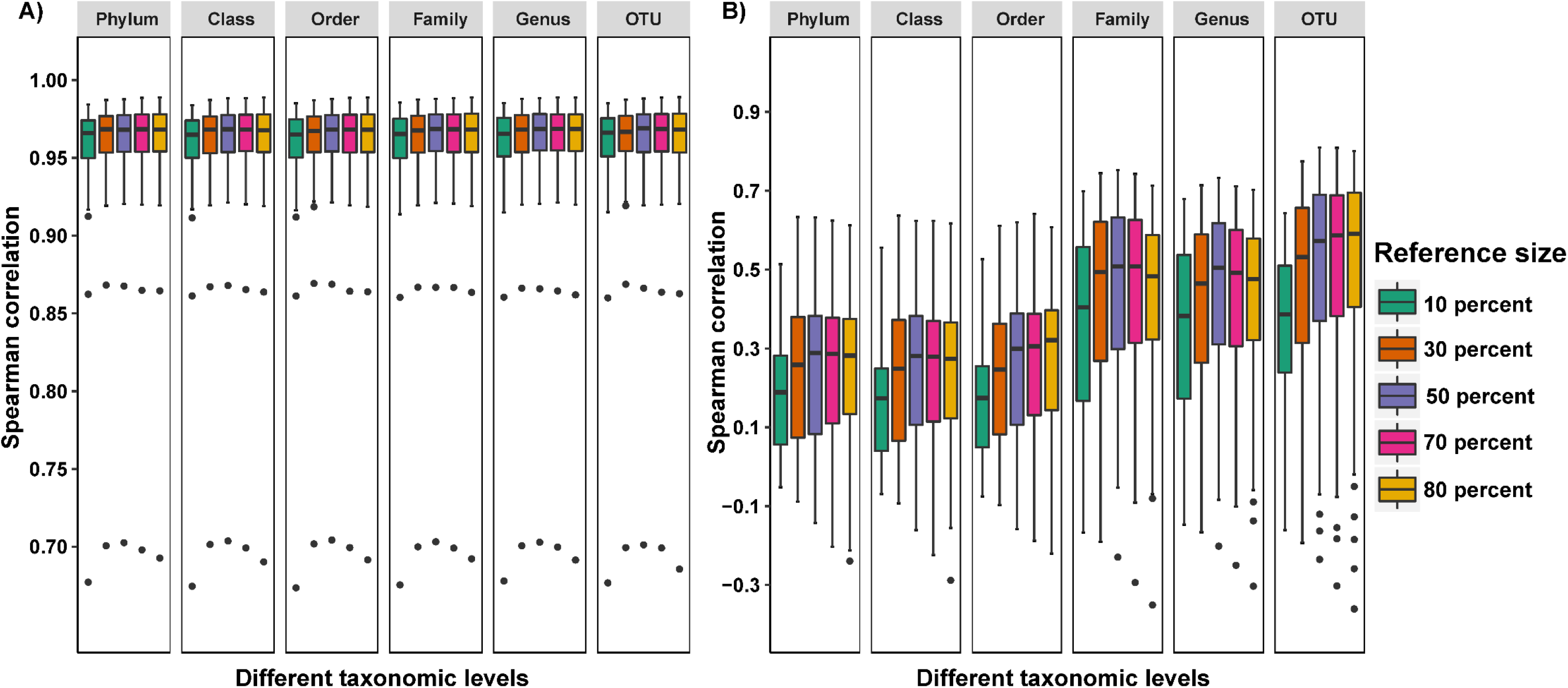
Comparison of sample and feature correlation using KEGG pathway abundance at different taxonomic levels and reference dataset size Boxplots showing the variation of **A)** Sample to sample correlations **B)** Feature to feature correlations obtained between the inferred KEGG pathway abundances and the mWGS functional profiles at all different reference sizes and taxonomic levels.

### IPCO outperforms PICRUSt, Tax4Fun and Piphillin in terms of both sample and feature correlation

To evaluate the functional inference of IPCO, we applied PICRUSt, Tax4Fun and Piphillin to the same datasets (**Table 1**) and KEGG pathway profiles were inferred. Spearman correlation was calculated for the inferred pathway profiles against its mWGS abundance both in terms of sample and feature correlation. IPCO outperformed PICRUSt, Tax4Fun and Piphillin in terms of sample correlation across all datasets (**Figure 3A**, **Supplementary table 3**). IPCO showed highest sample correlation with a narrow IQR range for stool and oral samples. Skin and nasal dataset showed lower sample correlation compared to stool and oral, however it was observed to be higher than what was observed using the other tools. Lowest sample correlation was observed using the environmental dataset but it was also higher than other tools for that site (**Supplementary table 3**).

**Figure 3.**
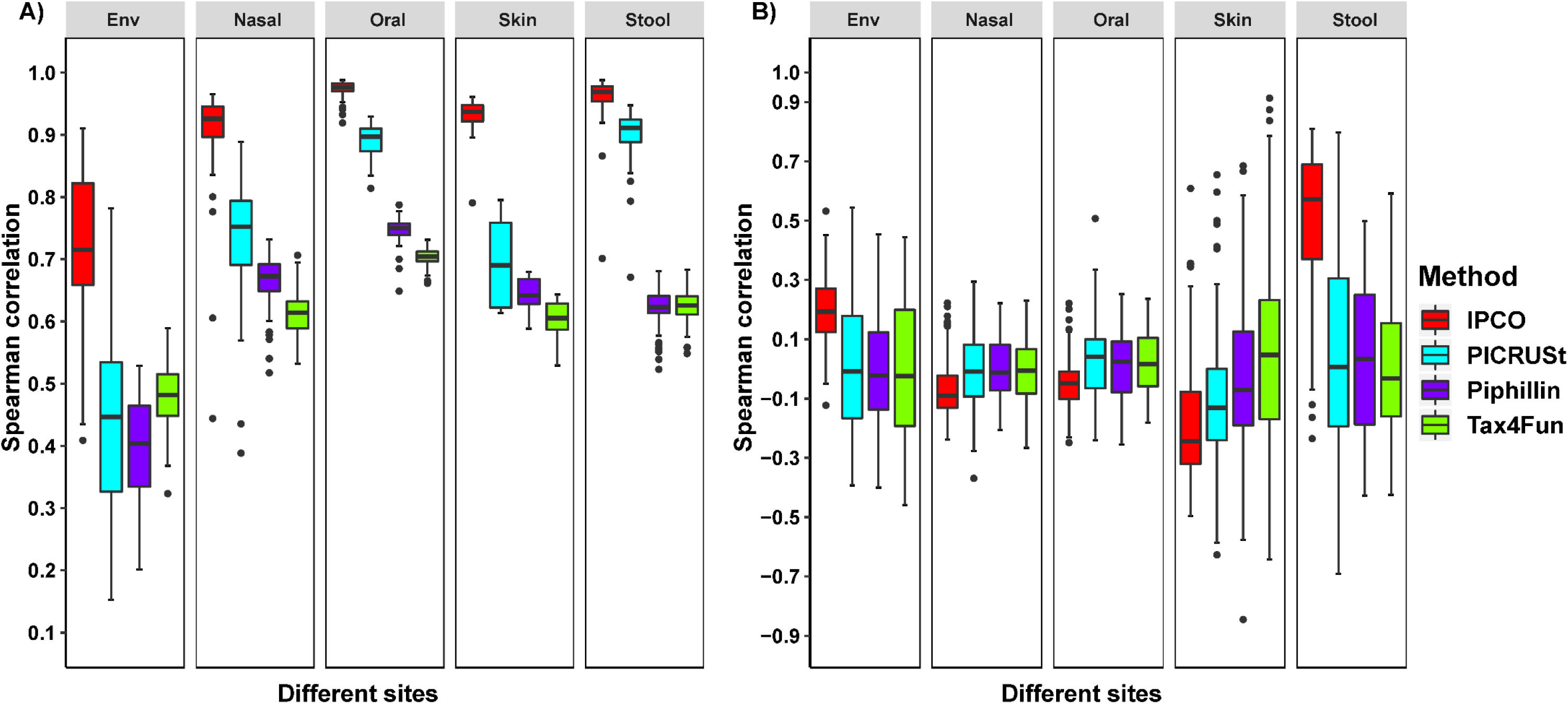
Sample and feature Spearman correlations of the inferred functional profiles with the mWGS functional profiles at different sites and using different methods Boxplots showing the comparison of **A)** Sample to sample correlations and **B)** Feature to feature correlations obtained between the inferred KEGG pathway abundance and the mWGS functional profiles at different sites using different methods.

Upon investing the feature correlations, it was observed that IPCO outperforms PICRUSt and Tax4Fun in stool and environmental dataset (**Figure 3B**). Nasal, oral and skin dataset revealed a lack of correlation using IPCO. It was noted that across all dataset, the median feature correlation for PICRUSt, Tax4Fun and Piphillin was close to zero (even with negative correlation values).

Sample and feature correlation based on KO abundances obtained from IPCO, PICRUSt, Tax4Fun and Piphillin were also calculated. IPCO was observed to outperform the other methods in terms of both sample correlation across all dataset as well as the feature correlations in the stool dataset.

### Lack of significant covariance between taxonomy and functional datasets results in lack of feature correlation in nasal, oral, skin and environmental dataset

To investigate the poor performance by IPCO on the other sites excluding stool, the co-variance between the taxonomic and functional dataset was calculated. Co-inertia analysis of the taxonomic profiles with its paired functional datasets across all sites reveal a lack of significant covariance between taxonomy and functional profiles in the nasal, oral, skin dataset (**Supplementary table 4**). Environmental dataset showed significant covariance if the mWGS derived taxonomy was used (while the 16S dataset did not co-vary significantly) (**Supplementary table 4**). This may in part explain the bad performance of IPCO on these datasets. Given the lack of covariance, further analysis was carried out only with stool dataset.

### Filtering by pathway coverage improves feature correlation in pathway datasets

Investigation of the effect of pathway coverage on the correlation values of the inferred pathways obtained using IPCO shows that coverage correlated well with pathway abundance and correlation values obtained with IPCO (**Figure 4A**). Based on this, for KEGG pathways, we identified that pathways below the mean coverage threshold of 0.01 were low correlated whereas pathways with a mean coverage between 0.01 and 0.1 showed a correlation value between 0.25 - 0.6. The highly correlated pathways have a mean coverage over 0.1% and showed high correction values between 0.6 -0.7(**Figure 4A**). In case of MetaCyc pathways, we observed similar results where pathways with coverage less than 0.41 (1^st^ quartile of mean coverage across samples) were low correlated pathways. Pathways whose mean coverage was between 1^st^ quartile (0.41) and median value (0.99) showed improved feature correlation for the inferred pathways whereas the best feature correlation (0.37 - 0.62) was observed for those pathways whose average pathway coverage was greater than its median value (**Figure 4B**). For both KEGG and MetaCyc, we were able to get high correlation values for more than 50% of the pathways based on the coverage filtering. The number of pathways retained in each of the coverage threshold, for both KEGG and MetaCyc are mentioned in **table 2**. For MetaCyc of 693 pathways are reported out of 712 because 19 pathways were detected only in one sample, hence during random bootstrap subsampling, these 19 pathways will present only in either reference or test datasets.

**Figure 4.**
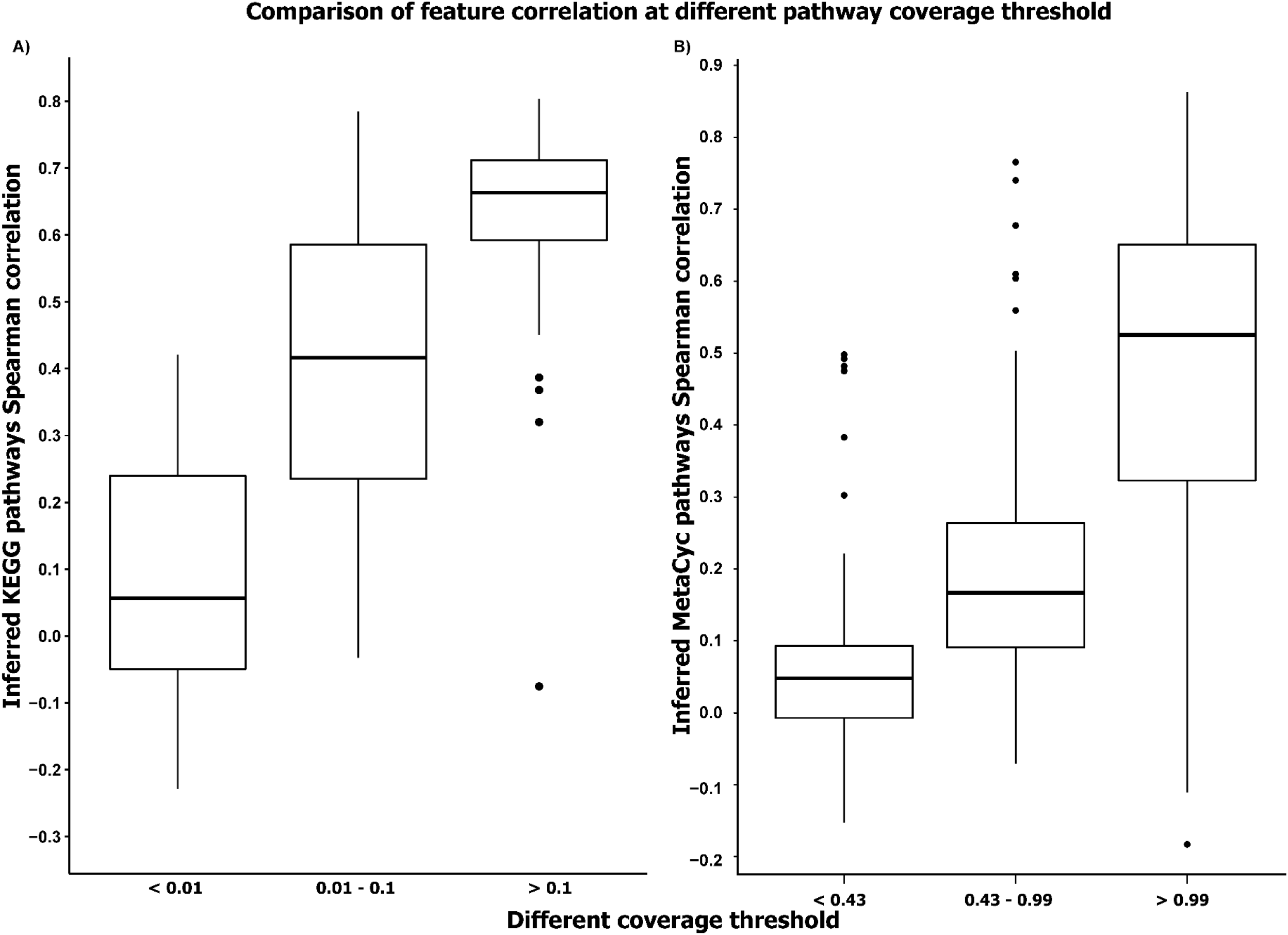
Comparison of feature correlations obtained using different pathway coverage thresholds for the KEGG and MetaCyc pathway classification schemes Boxplot showing the functional feature-to-feature correlations obtained using different coverage thresholds for the **A)** KEGG pathways and **B)** MetaCyc pathway schemes.

**Table 2.** Pathways retained at different coverage threshold

| **KEGG coverage threshold** | **Total** | **<0.01** | **0.01-0.1** | **>0.1** |
| --- | --- | --- | --- | --- |
| **Number of KEGG pathways** | 133 | 27 | 38 | 68 |
| **MetaCyc coverage threshold** | **Total** | **<0.43** | **0.43-0.99** | **>0.99** |
| **Number of MetaCyc pathways** | 693 | 173 | 166 | 354 |

Number of KEGG and MetaCyc pathways identified at different coverage threshold.

### Evaluation with an independent dataset shows that IPCO can infer the sample and features using any taxonomic dataset

Investigation of using mWGS species information as table L did not show significant difference as compared to using a 16S species or closed OTU level dataset as table L when working on an external 16S dataset to be inferred (**Supplementary figure 3**). This allowed consideration of more samples thus incorporation more functionality. **Table 3** describes the samples characteristics considered for this analysis.

**Table 3.**
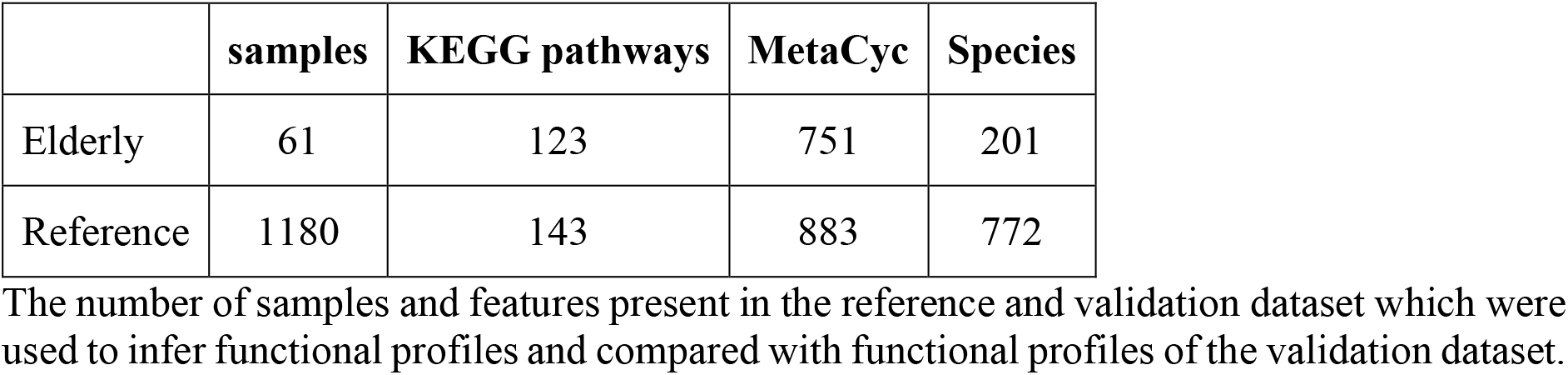
Number of samples, pathways and Species in the validation datasets

As it was observed that use of species levels datasets obtained from mWGS data was sufficient to inferred functionality on the elderly 16S dataset, the next part of the analysis was carried out by using the reference healthy functional and mWGS species profiles as reference in IPCO and the inferred functionality to the elderly 16S dataset.

### Inferred profiles from IPCO replicates mWGS functional profile associations with metabolites better than PICRUSt, Tax4Fun and Piphillin profiles

Correlation of the mWGS derived functionality (KEGG and MetaCyc) from elderly samples dataset to its paired bile acids and SCFA acid profiles have identified key pathways which directly correlate with the metabolite levels.

Investigation of the bile acid profiles showed that KEGG pathway “ko00121: Secondary bile acid biosynthesis” pathways significantly negatively correlates with primary bile acids (cholic acid and chenodeoxycholic acid) while “ko00790: Folate biosynthesis” known to promote bile acid levels [34] is observed to be significantly positively correlated with primary bile acids in elderly mWGS data.

With the secondary bile acids (lithocholic acid, dehydrocholic acid, 12-ketolithocholic acid, dehydrolithocholic acid, hyodeoxycholic acid and isolithocholic acid), it was observed that “ko00121: Secondary bile acid biosynthesis”, “ko00430: Taurine and hypotaurine metabolism”, “ko03070: Bacterial secretion system”, “ko05100: Bacterial invasion of epithelial cells” were all significantly positively correlated with secondary bile acid levels. This validates the bile acid profiles and the elderly mWGS functional dataset profiles in accordance to actual biochemical mechanism observed in a bacterial system.

The results of the inferred functional profiles obtained from all the tools shows that IPCO agrees best to the validation observed between the mWGS dataset and bile acid profiles (**Figure 5**, **Supplementary table 5**). It was observed that for primary bile acids only PICRUSt showed significant correlation with “ko00121: Secondary bile acid biosynthesis” however, the directionality was observed to be reversed. Tax4Fun and Piphillin did not show significant association their inferred “ko00121: Secondary bile acid biosynthesis” abundance and primary bile acids. Looking at the secondary bile acids, we observed that for 12-ketolithocholic acid did not show any significance with the inferred profiles obtained from all the tools. Lithocholic acid and KEGG pathways obtained from IPCO agreed with mWGS results and Tax4Fun despite showing significance showed opposite directionality. Dehydrolithocholic acid was significantly associated with KEGG pathways in PICRUSt and Piphillin. All associations, directionality and significance from all tools compared to mWGS results is highlighted in **figure 5**.

**Figure 5.**
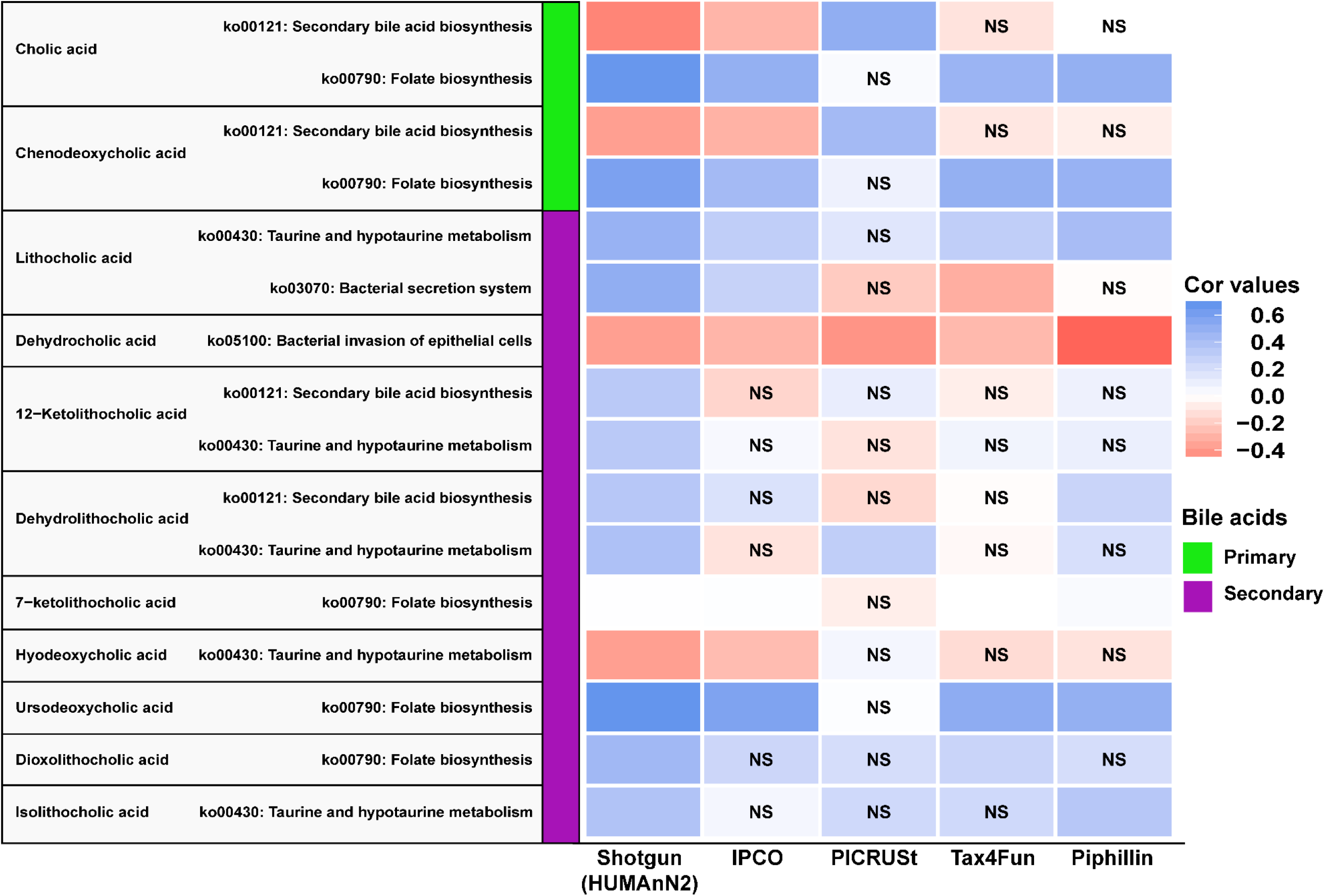
Correlation of the bile acids metabolite profiles obtained using the inferred KEGG pathway abundances from various methods Correlation of the inferred bile acid metabolite profile to paired mWGS KEGG pathways shows significant association (p-adjusted ≤ 0.1) with known pathways as shown in 1^st^ column. Directionally of association is shown by correlation values colour intensity. Pathways inferred from IPCO shows same directionality and significance (p-value ≤ 0.1) as observed with mWGS profiles for most cases. “**NS**” inside the cell represent non-significant (p-value > 0.1) associations.

The correlation between elderly KEGG pathway abundance and SCFA (butyrate and propionate) metabolite profiles observed non-significant correlation of butyrate and propionate levels with key KEGG pathways which included butanoate and propanoate metabolism pathways and protein and amino acid metabolism (Lysine, Glumatine). This lack of association was replicated in the inferred profiles obtained from IPCO and PICRUSt (**Supplementary table 6**). However, Tax4Fun and Piphillin showed significant association for the inferred KEGG pathways obtained using those two tools to both butyrate and propionate levels. These significant associations are considered false as they were not observed with the mWGS data. Piphillin reported the highest number of false positive pathways for butanoate. In case of propionate, IPCO also predicted two false positive pathways.

Similar results were observed when the MetaCyc pathway abundance dataset was used as reference. Correlation of the elderly mWGS MetaCyc pathways with bile acid profiles show significant correlation with “PWY-6518: glycocholate metabolism (bacteria)” and “1CMET2-PWY: N10-formyl-tetrahydrofolate biosynthesis”. These results were replicated with inferred MetaCyc pathway profiles obtained from IPCO (**Supplementary table 7**).

Investigation of the SCFAs (butyrate and propionate) levels with elderly mWGS MetaCyc pathways shows that butyrate shows no significant association with key butyrate MetaCyc pathways which was replicated with IPCO inferred functional dataset also. Propionate on the other hand showed significant association with few key MetaCyc amino acid metabolism pathways which couldn’t be replicated with IPCO (**Supplementary table 8**). The conflicting results observed with propionate could potentially be associated with the lower feature to feature correlation observed using MetaCyc as reference compared to high correlation obtained when using KEGG pathways dataset. As other tools do not report MetaCyc pathway profiles, this investigation could not be carried out with other tools.

## Discussion

We have developed IPCO, a novel tool which shows that prediction of functionality from 16S is not necessarily dependent on the phylogenetic information or their mapping to known reference taxa. The robustness between functionality and taxonomic profiles of microbiota is dependent up to a certain degree to not only the abundance of the various taxa but also the distribution of function across various microbes. However, alterations at taxonomic levels should also affect the overall functional potential at community level [35]. Using this concept, IPCO, is able to utilise the biologically and statistically significant covariance observed between the reference taxonomic and functional dataset and infer the functional capabilities of an external data to a great extent. IPCO was also implemented with both KEGG and MetaCyc pathway coverage as a threshold measure to eliminate pathways which do not correlate well. This allowed a selection of pathways which could be well predicted using IPCO.

The shortcoming of other tools is that given that most microbiome data consist of uncharacterised or partially characterised taxa, using a reference set of known functions from only a set of known taxa vastly limits the prediction of functionality for the whole amplicon dataset. Another major limitation that we have identified is that all these methods show a lack of feature to feature correlation i.e. their KEGG pathway abundance calculated from 16S datasets do not correlate well across samples obtained from a paired mWGS dataset. This is of concern as this potentially creates conflicting results when performing differential abundance analysis of the features and also show differing directionality when investigating functional profiles generated from 16S dataset against its paired metagenome dataset.

By studying various functionality associated with biologically relevant metabolites, e.g. in our case, bile acids and SCFAs, we have shown that the inferred functional capabilities obtained using IPCO, mimics the results of mWGS functional profiles and outperform other tools by a great extent. A comparison of such provides information relevant from a biological point of view as it is very important that the directionality of an observed significance is expected to be correct inference of the biological mechanism.

The filtering criteria in IPCO allows the users to select a set of functions with sufficient coverage to be inferred. This removes functions which may have been spuriously assigned due to the presence of only a small subset of genes/reactions. The reproducibility of results observed from both KEGG and MetaCyc shows that our method is independent of the database and can be easily implemented with custom datasets built from user’s internal data. Currently, IPCO can be implemented with any taxonomic level information and the taxonomic assignment can be done with any reference database as long as the taxa are present in the reference which acts as a mediator to co-vary the functional profiles with the 16S dataset.

Despite IPCO performing better than the other established tool, there are certain limitation to the tool. The contribution of a taxa for a functionality cannot be inferred using the IPCO algorithm. IPCO assigns a small pseudo value to each functionality due to the way the R’LQ algorithm calculates double co-inertia which makes the resulting inferred functionality non-zero abundant. To overcome this limitation, the low abundant functionality can easily be filtered by removing those functions whose average inferred abundance across samples is below a certain quantile determined by the user.

Overall, despite these limitations, IPCO provides a relatively better inference of functional potential than the available tools and can be easily implemented in R with very little requirements. With increasing mWGS data, more functionality can be easily added with deeper coverage allowing IPCO to be easily tuned to work at other sites. This would allow it to perform better at other sites which is not only allow better understanding of taxa function relationship but allow better functional prediction for external 16S dataset obtained from different sites.

## Notes

https://github.com/IPCO-Rlibrary/IPCO

