## Supplementary_figures for "IPCO: Inference of Pathways from Co-variance analysis"

**Supplementary figure 1** Sample-to-sample and feature-to-feature correlations between inferred and mWGS functional profiles obtained for MetaCyc pathway schemes using different transformations

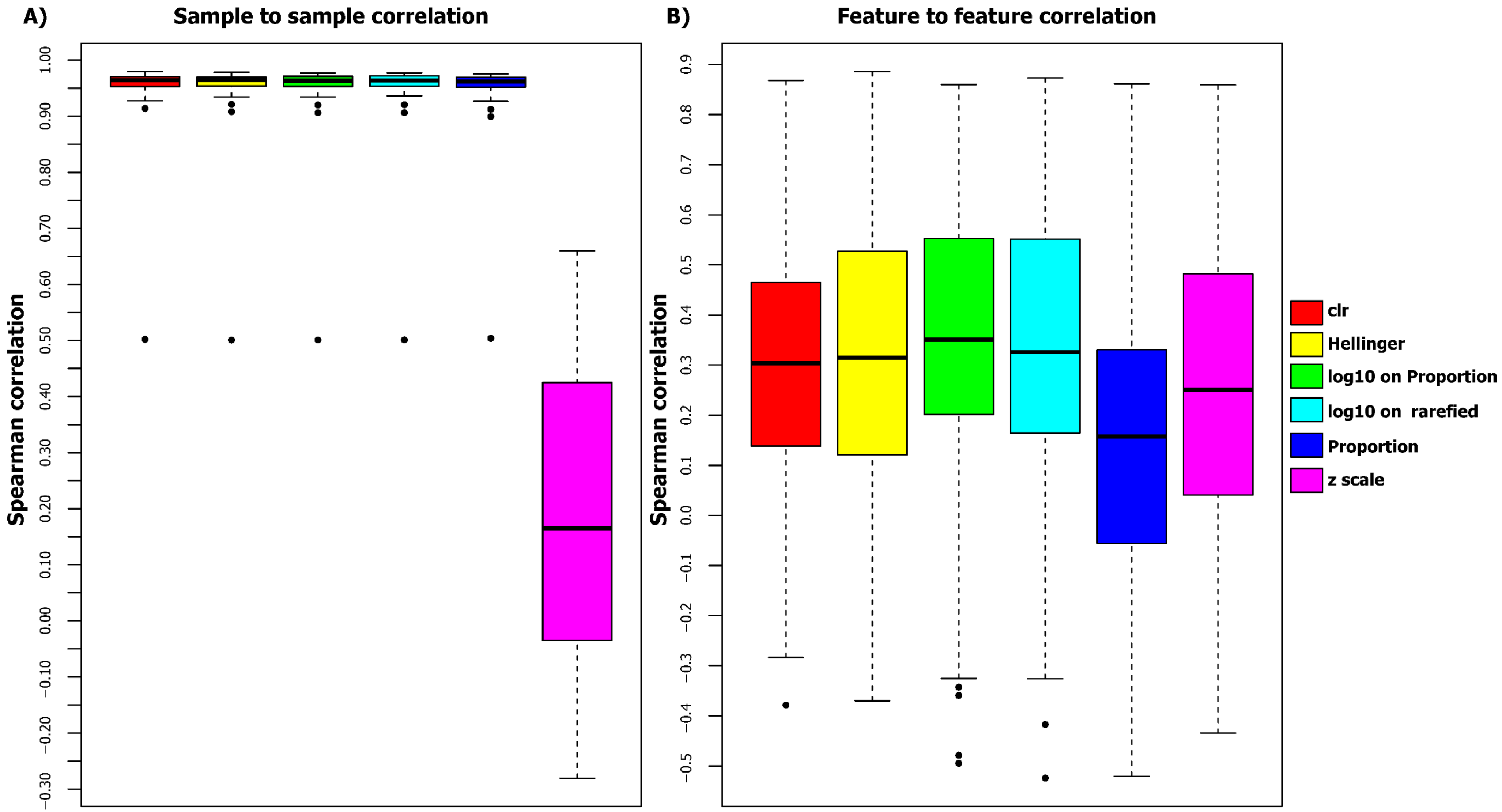


Supplementary figure 1 shows the effect of various transformation/normalisation methods based on preliminary analysis of IPCO's prediction using the MetaCyc functional profile datasets, in terms of **A)** sample to sample correlations and **B)** the correlation values by correlations all the IPCO inferred functional profiles to mWGS functional profiles

**Supplementary figure 2** Comparison of sample-to-sample and feature-to-feature correlations obtained between the inferred and the mWGS MetaCyc pathway abundances at different taxonomic levels and reference dataset size

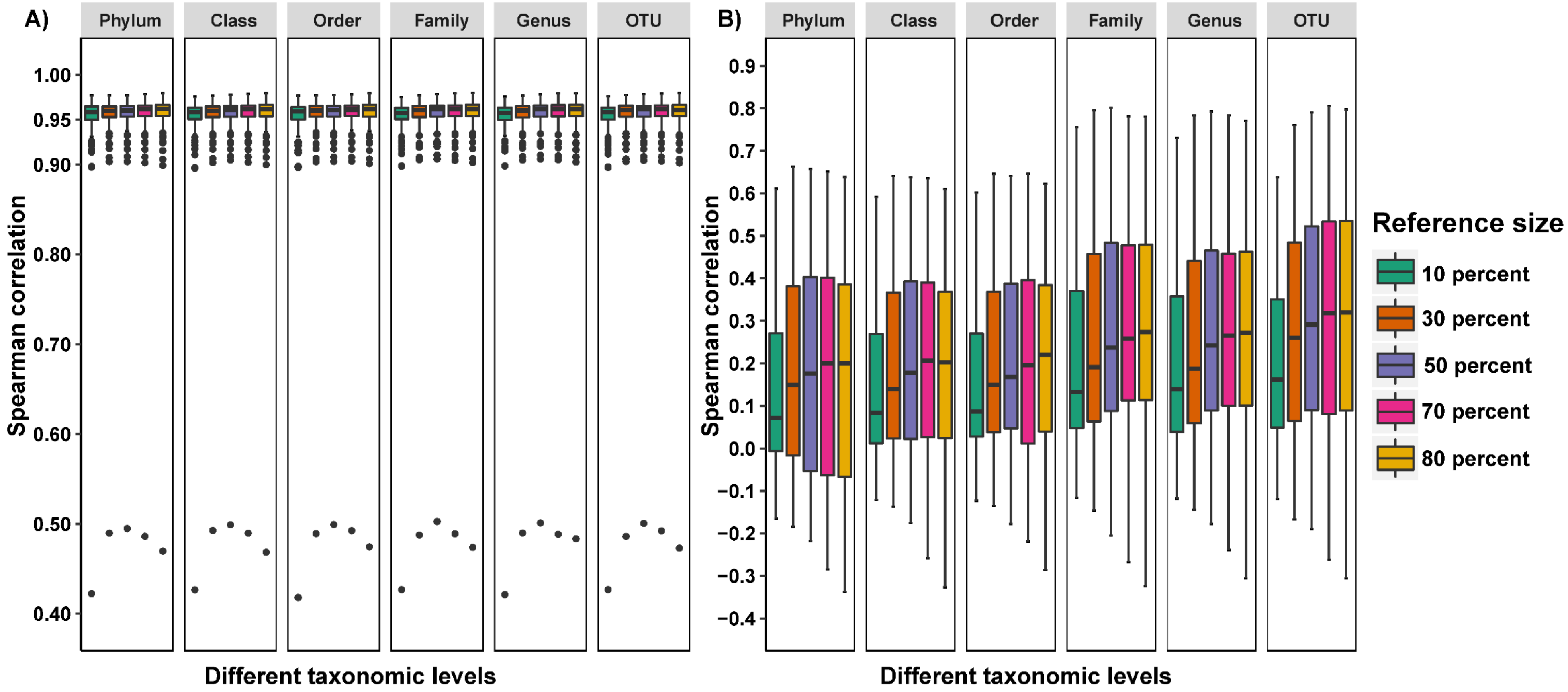


IPCO's prediction of MetaCyc pathways abundance using different reference dataset size and at different taxonomic levels shows **A)** high sample to sample correlation at all different reference size and taxonomic levels and **B)** shows the correlation values of features improve with larger reference and lowest taxonomic level

**Supplementary figure 3** Comparison of using 16S and mWGS taxonomic dataset as table L when inferring for external dataset

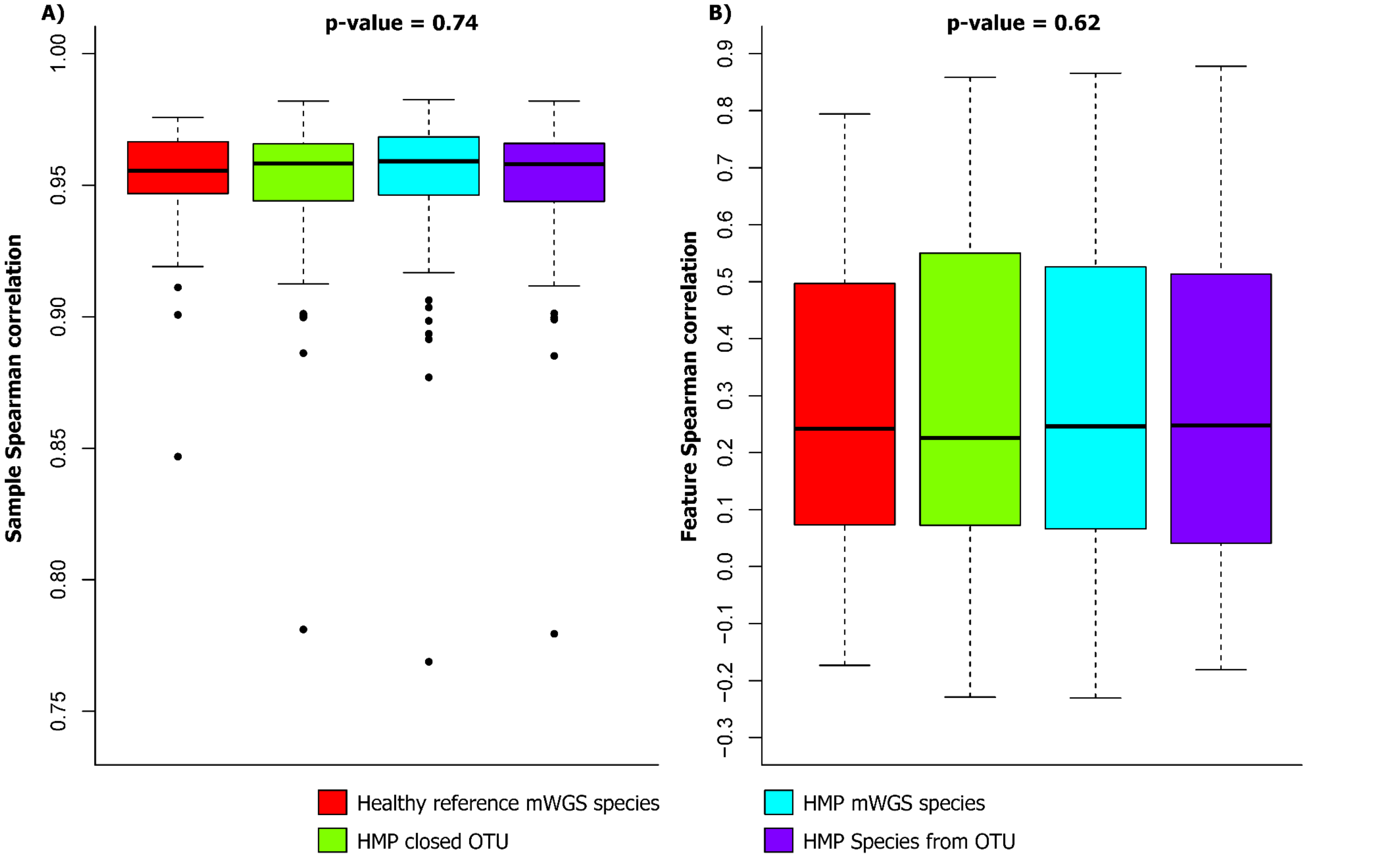


Supplementary figure 3 shows sample and feature correlation of inferred MetaCyc pathways of elderly 16S dataset obtained from using HMP 16S species and closed OTU, HMP mWGS species and healthy reference mWGS species as reference taxonomic dataset (table L). No significant change in observed sample (**A**) and feature (**B**) correlation for the external dataset when using the closed OTU level or species level dataset derived from either 16S representative sequences or mWGS taxonomy (**table L**). The external dataset (**table Q**) is collapsed to closed OTU level or species level derived from 16S representative sequences
