## Supplementary_tables for "IPCO: Inference of Pathways from Co-variance analysis"

**Supplementary table 1** Sample-to-sample and feature-to-feature covariance observed between mWGS and IPCO inferred functional profiles using different normalisation/transformation

| **Normalisation/transformation** | **Sample-to-sample covariance** | | **Feature-to-feature covariance** | |
| --- | --- | --- | --- | --- |
|  | **RV** | **P-value** | **RV** | **P-value** |
| Hellinger | 0.89 | 0.001 | 0.54 | 0.007 |
| Log10 on rarefied counts | 0.84 | 0.001 | 0.33 | 0.003 |
| Log10 on proportion | 0.84 | 0.001 | 0.31 | 0.002 |
| Proportion | 0.97 | 0.001 | 0.36 | 0.086 |
| z-scaling | 0.24 | 0.001 | 0.20 | 0.079 |
| clr transformation | 0.28 | 0.011 | 0.55 | 0.001 |

Comparison of the effect of different normalisation/transformation on the sample to sample and feature to feature co-inertia between IPCO inferred and mWGS MetaCyc pathabundance datasets. Co-variance is determined by RV coefficient. All observed co-variance were significant at nominal p-value ≤ 0.05 except for proportional and z-scaled features co-variance. Number of samples in reference dataset size is 70% more than the queried dataset.

**Supplementary table 2** Pairwise comparison of correlations observed using reference datasets of different size and taxonomic level

**A)** Significance (P-values) of the correlations between mWGS and IPCO-inferred KEGG pathway abundance profiles

| **Samples** | | | | | | |
| --- | --- | --- | --- | --- | --- | --- |
| **Reference size** | **Phylum** | **Class** | **Order** | **Family** | **Genus** | **OTU** |
| 10 - 30 | 2.6E-01 | 1.8E-01 | 3.1E-01 | 3.9E-01 | 3.0E-01 | 4.1E-01 |
| 10 - 50 | 2.1E-01 | 1.4E-01 | 2.1E-01 | 2.6E-01 | 2.6E-01 | 2.6E-01 |
| 10 - 70 | 2.2E-01 | 2.9E-01 | 5.0E-01 | 3.5E-01 | 6.7E-01 | 6.1E-01 |
| 10 - 80 | 4.1E-01 | 1.8E-01 | 2.5E-01 | 6.8E-01 | 3.6E-01 | 3.6E-01 |
| 30 - 50 | 5.6E-01 | 5.5E-01 | 5.0E-01 | 4.9E-01 | 5.7E-01 | 4.8E-01 |
| 30 - 70 | 5.5E-01 | 6.6E-01 | 6.2E-01 | 5.3E-01 | 7.4E-01 | 5.7E-01 |
| 30 - 80 | 6.3E-01 | 6.1E-01 | 5.1E-01 | 6.3E-01 | 6.5E-01 | 5.2E-01 |
| 50 - 70 | 4.8E-01 | 5.5E-01 | 5.7E-01 | 5.3E-01 | 5.9E-01 | 5.6E-01 |
| 50 - 80 | 5.2E-01 | 4.8E-01 | 5.1E-01 | 5.9E-01 | 4.9E-01 | 4.8E-01 |
| 70 - 80 | 4.9E-01 | 5.1E-01 | 5.0E-01 | 4.9E-01 | 5.4E-01 | 5.2E-01 |
| **Features** | | | | | | |
| **Reference size** | **Phylum** | **Class** | **Order** | **Family** | **Genus** | **OTU** |
| 10 - 30 | **1.2E-02** | **8.3E-04** | **7.2E-04** | **1.2E-02** | **3.4E-02** | **1.3E-05** |
| 10 - 50 | **3.7E-03** | **3.4E-05** | **2.3E-06** | **2.5E-03** | **2.6E-03** | **1.7E-08** |
| 10 - 70 | **2.9E-03** | **1.1E-04** | **6.2E-07** | **1.4E-03** | **3.1E-03** | **3.8E-09** |
| 10 - 80 | **4.2E-03** | **9.7E-05** | **3.0E-07** | **1.7E-02** | **1.3E-02** | **1.2E-09** |
| 30 - 50 | 4.7E-01 | 2.7E-01 | 1.2E-01 | 3.1E-01 | 2.1E-01 | 1.3E-01 |
| 30 - 70 | 4.3E-01 | 4.1E-01 | 7.2E-02 | 2.8E-01 | 2.6E-01 | 8.5E-02 |
| 30 - 80 | 4.6E-01 | 3.8E-01 | **4.8E-02** | 4.6E-01 | 3.6E-01 | 5.4E-02 |
| 50 - 70 | 4.8E-01 | 3.7E-01 | 4.1E-01 | 4.8E-01 | 4.0E-01 | 4.2E-01 |
| 50 - 80 | 4.8E-01 | 3.9E-01 | 3.5E-01 | 3.1E-01 | 3.0E-01 | 3.4E-01 |
| 70 - 80 | 5.2E-01 | 4.8E-01 | 4.0E-01 | 2.8E-01 | 3.6E-01 | 3.8E-01 |

Significance of correlation values of IPCO inferred samples and features obtained through pairwise comparison between different KEGG reference datasets size. Significance is determined by P-adjusted ≤ 0.05 and highlighted in red and bold

**B)** Significance (P-values) of the correlations between mWGS and IPCO-inferred MetaCyc pathway abundance profiles

| **Samples** | | | | | | |
| --- | --- | --- | --- | --- | --- | --- |
| **Reference size** | **Phylum** | **Class** | **Order** | **Family** | **Genus** | **OTU** |
| 10 - 30 | 3.8E-01 | 2.9E-01 | 4.8E-01 | 8.1E-02 | 2.2E-01 | 1.6E-01 |
| 10 - 50 | 4.4E-01 | 1.4E-01 | 4.2E-01 | **2.5E-02** | 7.4E-02 | 1.5E-01 |
| 10 - 70 | 4.4E-01 | 1.2E-01 | 4.0E-01 | **1.9E-02** | **4.9E-02** | 8.6E-02 |
| 10 - 80 | 4.1E-01 | 1.4E-01 | 4.0E-01 | **1.9E-02** | 5.9E-02 | 1.2E-01 |
| 30 - 50 | 3.8E-01 | 4.2E-01 | 4.4E-01 | 4.0E-01 | 3.7E-01 | 4.9E-01 |
| 30 - 70 | 3.8E-01 | 3.6E-01 | 4.2E-01 | 3.4E-01 | 2.7E-01 | 4.7E-01 |
| 30 - 80 | 4.4E-01 | 3.2E-01 | 3.8E-01 | 2.9E-01 | 2.5E-01 | 4.6E-01 |
| 50 - 70 | 4.2E-01 | 4.5E-01 | 4.0E-01 | 4.1E-01 | 4.1E-01 | 4.2E-01 |
| 50 - 80 | 4.2E-01 | 4.0E-01 | 4.5E-01 | 4.0E-01 | 3.8E-01 | 4.0E-01 |
| 70 - 80 | 3.9E-01 | 4.2E-01 | 4.6E-01 | 4.5E-01 | 4.3E-01 | 4.4E-01 |
| **Features** | | | | | | |
| **Reference size** | **Phylum** | **Class** | **Order** | **Family** | **Genus** | **OTU** |
| 10 - 30 | **2.2E-03** | **2.9E-05** | **2.3E-05** | **4.7E-05** | **3.3E-05** | **2.6E-08** |
| 10 - 50 | **1.9E-02** | **1.1E-06** | **5.9E-08** | **7.6E-09** | **3.9E-11** | **6.0E-16** |
| 10 - 70 | **1.7E-02** | **2.5E-07** | **9.6E-06** | **2.8E-11** | **4.0E-13** | **1.7E-16** |
| 10 - 80 | 7.1E-02 | **4.7E-06** | **5.8E-08** | **2.8E-11** | **3.3E-13** | **3.2E-17** |
| 30 - 50 | 2.8E-01 | 3.4E-01 | 1.6E-01 | 5.9E-02 | **9.4E-03** | **7.7E-03** |
| 30 - 70 | 2.9E-01 | 2.2E-01 | 4.5E-01 | **6.4E-03** | **1.2E-03** | **5.0E-03** |
| 30 - 80 | 1.1E-01 | 4.1E-01 | 1.6E-01 | **5.7E-03** | **1.0E-03** | **2.5E-03** |
| 50 - 70 | 4.6E-01 | 3.8E-01 | 1.8E-01 | 2.0E-01 | 2.7E-01 | 4.2E-01 |
| 50 - 80 | 2.8E-01 | 3.6E-01 | 4.5E-01 | 1.9E-01 | 2.6E-01 | 4.0E-01 |
| 70 - 80 | 2.8E-01 | 3.2E-01 | 1.7E-01 | 4.6E-01 | 4.5E-01 | 4.3E-01 |

Significance of correlation values of IPCO inferred samples and features obtained through pairwise comparison between different MetaCyc reference datasets size. Significance is determined by P-adjusted ≤ 0.05 and highlighted in red and bold

**Supplementary table 3** Comparison of sample-to-sample and feature-to-feature correlations between the inferred and the mWGS pathway profiles obtained using different methodologies for different sites

|  | **Samples** | | | | |
| --- | --- | --- | --- | --- | --- |
|  | **environmental** | **nasal** | **oral** | **skin** | **stool** |
| IPCO - PICRUSt | **2.3E-10** | **6.5E-08** | **1.7E-07** | **1.2E-02** | **8.9E-08** |
| IPCO - Piphillin | **1.7E-15** | **2.6E-18** | **1.1E-25** | **2.7E-03** | **1.7E-44** |
| IPCO - Tax4Fun | **2.2E-07** | **3.9E-35** | **7.3E-52** | **2.0E-05** | **2.7E-44** |
| PICRUSt - Piphillin | 5.4E-02 | **3.1E-04** | **5.6E-08** | 2.7E-01 | **1.4E-18** |
| PICRUSt - Tax4Fun | 1.1E-01 | **1.2E-12** | **6.2E-24** | **3.4E-02** | **1.6E-18** |
| Piphillin - Tax4Fun | **2.7E-03** | **1.4E-04** | **1.2E-06** | 9.9E-02 | 4.9E-01 |
|  | **Features** | | | | |
|  | **environmental** | **nasal** | **oral** | **skin** | **stool** |
| IPCO - PICRUSt | **2.4E-11** | **1.2E-06** | **2.9E-07** | **1.9E-03** | **9.4E-24** |
| IPCO - Piphillin | **1.3E-12** | **2.1E-07** | **5.0E-06** | **3.9E-08** | **8.5E-24** |
| IPCO - Tax4Fun | **3.1E-11** | **2.5E-07** | **2.0E-07** | **1.3E-12** | **4.6E-28** |
| PICRUSt - Piphillin | 3.2E-01 | 4.5E-01 | 3.2E-01 | **6.1E-03** | 5.0E-01 |
| PICRUSt - Tax4Fun | 5.0E-01 | 4.5E-01 | 4.3E-01 | **2.4E-05** | 3.0E-01 |
| Piphillin - Tax4Fun | 4.0E-01 | 4.2E-01 | 3.1E-01 | 5.2E-02 | 2.4E-01 |

Pairwise comparison of the correlation values from different methodology using dunn’s test. Significance is determined by P-adjusted ≤ 0.05 and highlighted in red and bold.

**Supplementary table 4** Co-variance observed between taxonomic and functional datasets from different sites

| **KEGG** | | | | |
| --- | --- | --- | --- | --- |
|  | **OTU and Pathway abundance** | | **mWGS species and Pathway abundance** | |
| **site** | **RV** | **pvalue** | **RV** | **pvalue** |
| **nasal** | 0.08 | 0.892 | 0.07 | 0.404 |
| **oral** | 0.14 | 0.845 | 0.12 | 0.366 |
| **skin** | 0.47 | 0.795 | 0.43 | 0.390 |
| **stool** | 0.35 | **0.001** | 0.35 | **0.001** |
| **environmental** | 0.17 | 0.188 | 0.23 | **0.009** |
| **MetaCyc** | | | | |
| **site** | **RV** | **pvalue** | **RV** | **pvalue** |
| **nasal** | 0.11 | 0.945 | 0.07 | 0.745 |
| **oral** | 0.14 | 0.911 | 0.13 | 0.233 |
| **skin** | 0.43 | 0.929 | 0.36 | 0.630 |
| **stool** | 0.41 | 0.001 | 0.47 | 0.001 |
| **environmental** | 0.19 | 0.146 | 0.22 | **0.012** |

Co-inertia of samples and pathways between the inferred KEGG/MetaCyc profiles with its paired mWGS pathabundance. Degree of co-variance is determined by RV coefficient and significance is determined by nominal p-value ≤ 0.05 and highlighted in red and bold

**Supplementary table 5** Correlation between the paired bile acid profiles and the different KEGG pathways obtained using mWGS and those inferred using different methods

| **Spearman correlation values** | | | | | | |
| --- | --- | --- | --- | --- | --- | --- |
| **Bile_acids** | **KEGG pathways** | **Shotgun** | **IPCO** | **PICRUSt** | **Tax4Fun** | **Piphillin** |
| **cholic_acid** | ko00121: Secondary bile acid biosynthesis | -0.34 | -0.21 | 0.37 | -0.08 | 0.00 |
| **cholic_acid** | ko00790: Folate biosynthesis | 0.48 | 0.36 | 0.02 | 0.34 | 0.36 |
| **Chenodeoxycholic_acid** | ko00121: Secondary bile acid biosynthesis | -0.27 | -0.22 | 0.31 | -0.07 | -0.05 |
| **Chenodeoxycholic_acid** | ko00790: Folate biosynthesis | 0.43 | 0.31 | 0.06 | 0.36 | 0.35 |
| **Lithocholic_acid** | ko00430: Taurine and hypotaurine metabolism | 0.35 | 0.22 | 0.11 | 0.22 | 0.3 |
| **Lithocholic_acid** | ko03070: Bacterial secretion system | 0.37 | 0.2 | -0.15 | -0.23 | -0.01 |
| **Dehydrocholic_acid** | ko05100: Bacterial invasion of epithelial cells | -0.27 | -0.21 | -0.30 | -0.20 | -0.42 |
| **12-Ketolithocholic_acid** | ko00121: Secondary bile acid biosynthesis | 0.24 | -0.12 | 0.08 | -0.05 | 0.06 |
| **12-Ketolithocholic_acid** | ko00430: Taurine and hypotaurine metabolism | 0.24 | 0.03 | -0.08 | 0.05 | 0.07 |
| **dehydrolithocholic_acid** | ko00121: Secondary bile acid biosynthesis | 0.25 | 0.13 | -0.11 | -0.01 | 0.19 |
| **dehydrolithocholic_acid** | ko00430: Taurine and hypotaurine metabolism | 0.28 | -0.08 | 0.22 | -0.02 | 0.14 |
| **7-ketolithocholic_acid** | ko00790: Folate biosynthesis | 0.003 | 0.006 | -0.05 | 0.001 | 0.021 |
| **Hyodeoxycholic_acid** | ko00430: Taurine and hypotaurine metabolism | -0.27 | -0.19 | 0.04 | -0.10 | -0.08 |
| **Ursodeoxycholic_acid** | ko00790: Folate biosynthesis | 0.49 | 0.42 | 0.01 | 0.38 | 0.36 |
| **Dioxolithocholic_acid** | ko00790: Folate biosynthesis | 0.32 | 0.18 | 0.15 | 0.19 | 0.15 |
| **Isolithocholic_acid** | ko00430: Taurine and hypotaurine metabolism | 0.27 | 0.04 | 0.17 | 0.17 | 0.25 |
| **P-values of Spearman correlation** | | | | | | |
| **cholic_acid** | ko00121: Secondary bile acid biosynthesis | **0.014** | 0.064 | **0.001** | 0.497 | 0.971 |
| **cholic_acid** | ko00790: Folate biosynthesis | **0.001** | **0.001** | 0.865 | **0.002** | **0.001** |
| **Chenodeoxycholic_acid** | ko00121: Secondary bile acid biosynthesis | **0.052** | **0.049** | **0.005** | 0.521 | 0.638 |
| **Chenodeoxycholic_acid** | ko00790: Folate biosynthesis | **0.002** | **0.006** | 0.605 | **0.001** | **0.001** |
| **Lithocholic_acid** | ko00430: Taurine and hypotaurine metabolism | **0.013** | **0.049** | 0.340 | 0.051 | **0.007** |
| **Lithocholic_acid** | ko03070: Bacterial secretion system | **0.008** | 0.08 | 0.180 | **0.04** | 0.927 |
| **Dehydrocholic_acid** | ko05100: Bacterial invasion of epithelial cells | **0.053** | 0.059 | **0.006** | 0.084 | **0.000** |
| **12-Ketolithocholic_acid** | ko00121: Secondary bile acid biosynthesis | **0.099** | 0.293 | 0.492 | 0.68 | 0.573 |
| **12-Ketolithocholic_acid** | ko00430: Taurine and hypotaurine metabolism | **0.085** | 0.809 | 0.472 | 0.647 | 0.541 |
| **dehydrolithocholic_acid** | ko00121: Secondary bile acid biosynthesis | **0.072** | 0.251 | 0.326 | 0.897 | 0.089 |
| **dehydrolithocholic_acid** | ko00430: Taurine and hypotaurine metabolism | **0.048** | 0.462 | 0.056 | 0.842 | 0.207 |
| **7-ketolithocholic_acid** | ko00790: Folate biosynthesis | **0.003** | **0.006** | 0.673 | **0.001** | **0.021** |
| **Hyodeoxycholic_acid** | ko00430: Taurine and hypotaurine metabolism | **0.051** | 0.089 | 0.739 | 0.360 | 0.479 |
| **Ursodeoxycholic_acid** | ko00790: Folate biosynthesis | **0.001** | **0.0001** | 0.943 | **0.001** | **0.001** |
| **Dioxolithocholic_acid** | ko00790: Folate biosynthesis | **0.02** | 0.123 | **0.191** | 0.098 | 0.178 |
| **Isolithocholic_acid** | ko00430: Taurine and hypotaurine metabolism | **0.055** | 0.757 | 0.131 | 0.138 | **0.025** |

Correlation between mWGS KEGG pathways profiles with its paired bile acid metabolites levels and replication of correlation between inferred profiles from different methodology with bile acids levels. Significance for the mWGS profile is determined by p-adjusted ≤ 0.1 and for the different methodology by nominal p-value ≤ 0.1 and highlighted in red and bold. Directionality and strength of correlation is determined by spearman correlation estimate.

**Supplementary table 6** Correlation of mWGS and inferred KEGG pathways to its paired butyrate and propionate levels

|  |  |  | **P-values** | | | | |
| --- | --- | --- | --- | --- | --- | --- | --- |
|  | | | **shotgun** | **IPCO** | **PICRUSt** | **Tax4Fun** | **Piphillin** |
| **Butyrate** | ko00250: | Alanine, aspartate and glutamate metabolism | 0.59 | 0.83 | 0.58 | 0.57 | **0.06** |
|  | ko00300: | Lysine biosynthesis | 0.13 | 0.76 | 0.59 | 0.79 | **0.04** |
|  | ko00310: | Lysine degradation | 0.44 | 0.24 | 0.76 | 0.74 | 0.39 |
|  | ko00471: | D-Glutamine and D-glutamate metabolism | 0.48 | 0.83 | 0.86 | **0.05** | 0.31 |
|  | ko00650: | Butanoate metabolism | 0.29 | 0.32 | 0.46 | 0.68 | **0.09** |
|  | ko04974: | Protein digestion and absorption | 0.56 | 0.87 | 0.80 | 0.43 | 0.70 |
| **Propionate** | ko00250: | Alanine, aspartate and glutamate metabolism | 0.52 | 0.90 | 0.55 | 0.37 | 0.19 |
|  | ko00260: | Glycine, serine and threonine metabolism | 0.35 | 0.95 | 0.93 | 0.50 | 0.14 |
|  | ko00270: | Cysteine and methionine metabolism | 0.31 | 0.12 | 0.77 | 0.23 | 0.57 |
|  | ko00340: | Histidine metabolism | 0.39 | 0.89 | 0.21 | 0.99 | **0.01** |
|  | ko00471: | D-Glutamine and D-glutamate metabolism | 0.47 | 0.43 | 0.97 | **0.01** | 0.64 |
|  | ko00473: | D-Alanine metabolism | 0.72 | **0.01** | 0.96 | 0.22 | 0.93 |
|  | ko00640: | Propanoate metabolism | 0.35 | **0.07** | 0.32 | 0.13 | 0.23 |
|  | ko04974: | Protein digestion and absorption | 0.68 | 1.00 | 0.39 | 0.13 | 0.14 |

Correlation between mWGS KEGG pathways profiles with its paired butyrate and propionate metabolites levels and replication of correlation between inferred profiles from different methodology with butyrate and propionate levels. Significance for the mWGS profile and for the different methodology by nominal p-value ≤ 0.1 and highlighted in red and bold. mWGS profiles were not significantly correlated with butyrate and propionate levels.

**Supplementary table 7** Correlation between mWGS and inferred MetaCyc pathway pathway abundance and their paired bile acid profiles

|  |  | **Shotgun** | | **IPCO** | |
| --- | --- | --- | --- | --- | --- |
| **Bile_acids** | **MetaCyc pathways** | **cor value** | **padj** | **cor value** | **pval** |
| **cholic acid** | 1CMET2-PWY: N10-formyl-tetrahydrofolate biosynthesis | 0.49 | **0.001** | 0.33 | **0.003** |
| **cholic acid** | PWY-6518: glycocholate metabolism (bacteria) | 0.35 | **0.034** | 0.37 | **0.001** |
| **Chenodeoxycholic acid** | 1CMET2-PWY: N10-formyl-tetrahydrofolate biosynthesis | 0.45 | **0.004** | 0.29 | **0.010** |
| **Chenodeoxycholic acid** | PWY-6518: glycocholate metabolism (bacteria) | 0.31 | **0.061** | 0.33 | **0.003** |
| **Deoxycholic acid** | 1CMET2-PWY: N10-formyl-tetrahydrofolate biosynthesis | 0.38 | **0.015** | 0.25 | **0.024** |
| **Deoxycholic acid** | PWY-6518: glycocholate metabolism (bacteria) | 0.36 | **0.026** | 0.25 | **0.025** |
| **lithocholic acid** | 1CMET2-PWY: N10-formyl-tetrahydrofolate biosynthesis | 0.32 | **0.058** | 0.21 | **0.063** |
| **lithocholic acid** | PWY-6518: glycocholate metabolism (bacteria) | 0.30 | **0.082** | 0.26 | **0.020** |
| **Dehydrocholic acid** | 1CMET2-PWY: N10-formyl-tetrahydrofolate biosynthesis | 0.43 | **0.005** | 0.29 | **0.010** |
| **7-ketolithocholic acid** | 1CMET2-PWY: N10-formyl-tetrahydrofolate biosynthesis | 0.40 | **0.096** | 0.27 | **0.018** |
| **7-ketolithocholic acid** | PWY-6518: glycocholate metabolism (bacteria) | 0.29 | **0.009** | 0.33 | **0.003** |
| **Ursodeoxycholic_acid** | 1CMET2-PWY: N10-formyl-tetrahydrofolate biosynthesis | 0.50 | **0.001** | 0.40 | **0.0003** |
| **Ursodeoxycholic_acid** | PWY-6518: glycocholate metabolism (bacteria) | 0.29 | **0.087** | 0.38 | **0.001** |
| **Dioxolithocholic_acid** | 1CMET2-PWY: N10-formyl-tetrahydrofolate biosynthesis | 0.34 | **0.041** | 0.15 | 0.183 |

Correlation between mWGS MetaCyc pathways profiles with its paired bile acid metabolites levels and replication of correlation between IPCO inferred profiles with bile acids levels. Significance for the mWGS profile is determined by p-adjusted ≤ 0.1 and for IPCO profiles by nominal p-value ≤ 0.1 and highlighted in red and bold. Directionality and strength of correlation is determined by spearman correlation estimate.

**Supplementary table 8** P-values of correlation from mWGS and inferred MetaCyc pathways to its paired butyrate and propionate levels

|  |  | **shotgun** | **IPCO** |
| --- | --- | --- | --- |
| **Butyrate** | ARGDEG-PWY: superpathway of L-arginine, putrescine, and 4-aminobutanoate degradation | 0.80 | 0.93 |
|  | CENTFERM-PWY: pyruvate fermentation to butanoate | 0.65 | 0.30 |
|  | DAPLYSINESYN-PWY: L-lysine biosynthesis I | 0.791 | 0.19 |
|  | P162-PWY: L-glutamate degradation V (via hydroxyglutarate) | 0.31 | 0.77 |
|  | P163-PWY: L-lysine fermentation to acetate and butanoate | 0.69 | 0.27 |
|  | P4-PWY: superpathway of L-lysine, L-threonine and L-methionine biosynthesis I | 0.842 | **0.09** |
|  | PWY-2941: L-lysine biosynthesis II | 0.435 | 0.25 |
|  | PWY-2942: L-lysine biosynthesis III | **0.004** | **0.08** |
|  | PWY-5022: 4-aminobutanoate degradation V | 0.62 | 0.18 |
|  | PWY-5097: L-lysine biosynthesis VI | **0.002** | 0.19 |
|  | PWY-5100: pyruvate fermentation to acetate and lactate II | 0.88 | 0.49 |
|  | PWY-5177: glutaryl-CoA degradation | 0.90 | 0.64 |
|  | PWY-5505: L-glutamate and L-glutamine biosynthesis | 0.39 | 0.20 |
|  | PWY-5676: acetyl-CoA fermentation to butanoate II | 0.47 | 0.89 |
|  | PWY-5677: succinate fermentation to butanoate | 0.11 | 0.59 |
|  | PWY-6590: superpathway of Clostridium acetobutylicum acidogenic fermentation | 0.64 | 0.33 |
| **Propionate** | ASPASN-PWY: superpathway of L-aspartate and L-asparagine biosynthesis | **0.08** | 0.41 |
|  | HISDEG-PWY: L-histidine degradation I | 0.92 | **0.05** |
|  | HISTSYN-PWY: L-histidine biosynthesis | 0.12 | **0.001** |
|  | HOMOSER-METSYN-PWY: L-methionine biosynthesis I | 0.89 | 0.28 |
|  | HSERMETANA-PWY: L-methionine biosynthesis III | 0.37 | **0.04** |
|  | MET-SAM-PWY: superpathway of S-adenosyl-L-methionine biosynthesis | 0.88 | **0.01** |
|  | METSYN-PWY: L-homoserine and L-methionine biosynthesis | 0.70 | **0.01** |
|  | P108-PWY: pyruvate fermentation to propanoate I | 0.84 | 0.95 |
|  | P162-PWY: L-glutamate degradation V (via hydroxyglutarate) | 0.63 | 0.45 |
|  | P4-PWY: superpathway of L-lysine, L-threonine and L-methionine biosynthesis I | 0.98 | **0.01** |
|  | PROPFERM-PWY: L-alanine fermentation to propanoate and acetate | 0.45 | 0.89 |
|  | PWY0-1061: superpathway of L-alanine biosynthesis | 0.19 | 0.14 |
|  | PWY0-781: aspartate superpathway | 0.93 | **0.01** |
|  | PWY-5028: L-histidine degradation II | 0.29 | 0.96 |
|  | PWY-5088: L-glutamate degradation VIII (to propanoate) | **0.06** | 0.23 |
|  | PWY-5100: pyruvate fermentation to acetate and lactate II | 0.84 | 0.15 |
|  | PWY-5345: superpathway of L-methionine biosynthesis (by sulfhydrylation) | 0.56 | 0.11 |
|  | PWY-5347: superpathway of L-methionine biosynthesis (transsulfuration) | 0.93 | **0.02** |
|  | PWY-5505: L-glutamate and L-glutamine biosynthesis | 0.37 | **0.08** |
|  | PWY-6151: S-adenosyl-L-methionine cycle I | **0.01** | 0.51 |
|  | PWY-6628: superpathway of L-phenylalanine biosynthesis | 0.19 | 0.77 |
|  | PWY-7528: L-methionine salvage cycle I (bacteria and plants) | 0.50 | **0.10** |
|  | SER-GLYSYN-PWY: superpathway of L-serine and glycine biosynthesis I | 0.85 | 0.92 |
|  | THREOCAT-PWY: superpathway of L-threonine metabolism | 0.28 | 0.40 |
|  | THRESYN-PWY: superpathway of L-threonine biosynthesis | 0.46 | 0.34 |

Correlation between mWGS MetaCyc pathways profiles with its paired butyrate and propionate metabolites levels and replication of correlation between IPCO inferred profiles with butyrate and propionate levels. Significance for the mWGS profile and IPCO profiles is determined by nominal p-value ≤ 0.1 and highlighted in red and bold. mWGS profiles were not significantly correlated with butyrate and only three pathways with propionate levels.
